# Structures of filaments from Pick’s disease reveal a novel tau protein fold

**DOI:** 10.1101/302216

**Authors:** Benjamin Falcon, Wenjuan Zhang, Alexey G. Murzin, Garib Murshudov, Holly J. Garringer, Ruben Vidal, R. Anthony Crowther, Bernardino Ghetti, Sjors H.W. Scheres, Michel Goedert

## Abstract

The ordered assembly of tau protein into abnormal filamentous inclusions underlies many human neurodegenerative diseases^1^. Tau assemblies appear to spread through specific neural networks in each disease^2^, with short filaments having the greatest seeding activity^3^. The abundance of tau inclusions strongly correlates with disease symptoms^4^. Six tau isoforms are expressed in normal adult human brain - three isoforms with four microtubule-binding repeats each (4R tau) and three isoforms lacking the second repeat (3R tau)^1^. In various diseases, tau filaments can be composed of either 3R tau or 4R tau, or of both 3R and 4R tau. They have distinct cellular and neuroanatomical distributions^5^, with morphological and biochemical differences suggesting that they may be able to adopt disease-specific molecular conformations^6,7^. Such conformers may give rise to different neuropathological phenotypes^8,9^, reminiscent of prion strains^10^. However, the underlying structures are not known. Using electron cryo-microscopy (cryo-EM), we recently reported the structures of tau filaments from Alzheimer’s disease, which contain both 3R and 4R tau^11^. Here we have determined the structures of tau filaments from Pick’s disease, a neurodegenerative disorder characterised by frontotemporal dementia. They consist of residues K_254_-F_378_ of 3R tau, which are folded differently when compared to tau in Alzheimer’s disease filaments, establishing the existence of conformers of assembled tau. The Pick fold explains the selective incorporation of 3R tau in Pick bodies and the differences in phosphorylation relative to the tau filaments of Alzheimer’s disease. Our findings show how tau can adopt distinct folds in human brain in different diseases, an essential step for understanding the formation and propagation of molecular conformers.

---

We used cryo-EM to image tau filaments extracted from the frontotemporal cortex of a patient who had a 7 year history of behavioural-variant frontotemporal dementia. Neuropathological examination revealed severe frontotemporal lobar degeneration, with abundant Pick bodies composed of 3R tau filaments, without phosphorylation of S_262_ (Fig. 1a-d, Extended Data Fig. 1, Extended Data Table l)^12-17^. As in Alzheimer’s disease^18^, a fuzzy coat composed of the disordered N- and C-terminal regions of tau surrounded the filament cores and was removed by mild pronase treatment (Fig. 1e and Extended Data Fig. 1).

**Figure 1:**
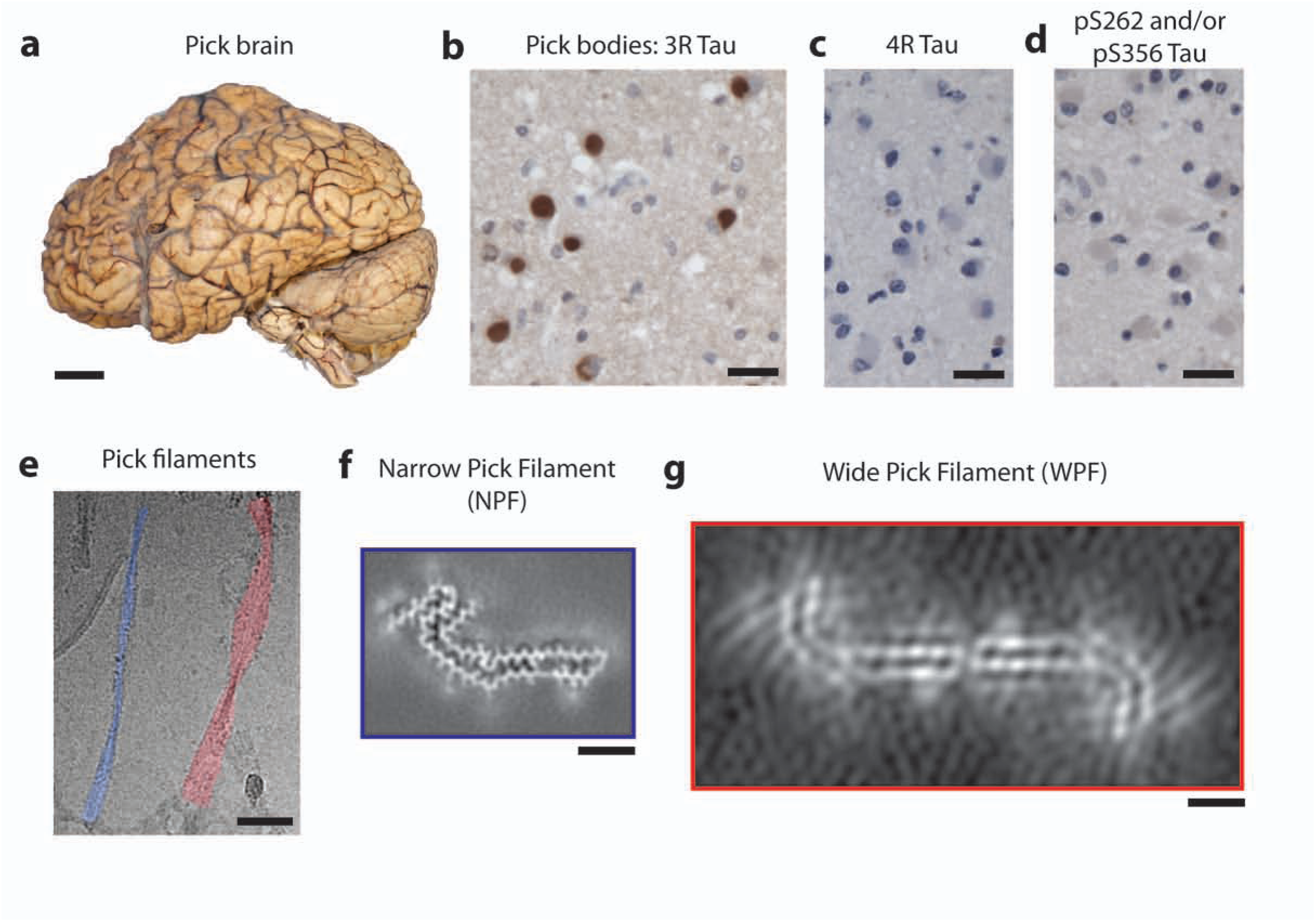
Filamentous tau pathology of Pick’s disease. **a.** The brain used in this study showed atrophy of anterior frontal and temporal lobes of the cerebral cortex. Grey matter from frontotemporal cortex was used for cryo-EM. Scale bar, 5 cm **b-d.** Staining of Pick bodies in frontotemporal cortex by RD3 (3RTau; brown) (**b.**), but not by anti-4R (4R Tau) (**c**.) or 12E8 (pS262 tau and/or pS356 tau) (**d**.). Nuclei were counterstained blue. Scale bars, 20 μm. **e.** Cryo-electron micrograph of extracted tau filaments, in which narrow (NPFs; false coloured blue) and wide (WPFs; false coloured red) Pick filaments could be distinguished. Scale bar, 500 Å **f.** Unsharpened cryo-EM density of NPF. Scale bar, 25 Å. **g.** Unsharpened cryo-EM density of WPF. Scale bar, 25 Å.

Narrow (93%) and wide (7%) filaments could be distinguished (Fig. 1e). The narrow filaments have previously been described as straight^19-21^, but they do have a helical twist with a cross-over distance of ~1000 Å and a projected width varying from approximately 50 to 150 Å. The wide filaments have a similar cross-over distance, but their width varies from approximately 50 to 300 Å. We named them narrow and wide Pick filaments (NPFs and WPFs). Their morphologies and relative abundance match those reported in cortical biopsies from Pick’s disease brain ^21^.

Using helical reconstruction in RELION^22^, we determined a 3.2 Å resolution map of the ordered core of NPFs, in which side-chain densities were well resolved and β-strands were clearly separated along the helical axis (Fig. 1f and Extended Data Fig. 2). We also determined an 8 Å resolution map of WPFs, which showed well-separated β-sheets perpendicular to the helical axis, but no separation of β-strands along the helical axis (Fig. 1g and Extended Data Fig. 3). NPFs are composed of a single protofilament with an elongated structure that is markedly different from the C-shaped protofilament of Alzheimer’s disease paired helical and straight filaments (PHFs and SFs)^11,23^. WPFs are formed by the association of two NPF protofilaments at their distal tips. In support, we observed WPFs where one protofilament had been lost in some parts (Extended Data Fig. 3). Our results reveal that the tau filaments of Pick’s disease adopt a single fold that is different from that of the tau filaments of Alzheimer’s disease.

The high-resolution NPF map allowed us to build an atomic model of the Pick tau filament fold, which consists of residues K_254_-F_378_ of 3R tau (in the numbering of the 441 amino acid human tau isoform) (Fig. 2). There are nine β-strands (β1-9) arranged into four cross-β packing stacks and connected by turns and arcs (Fig. 3a,b). R1 provides two strands, *β*1 and *β*2, and R3 and R4 provide three strands each, *β*3-5 and *β*6-8, respectively. These pack together in a hairpin-like fashion: *β*1 against *β*8, *β*2 against *β*7, *β*3 against *β*6 and *β*4 against *β*5. The final strand, *β*9, is formed from 9 amino acids after R4 and packs against the opposite side of *β*8. Only the interface between β3 and β6 is entirely hydrophobic; the other cross-β packing interfaces are composed of both non-polar and polar side-chains.

**Figure 2:**
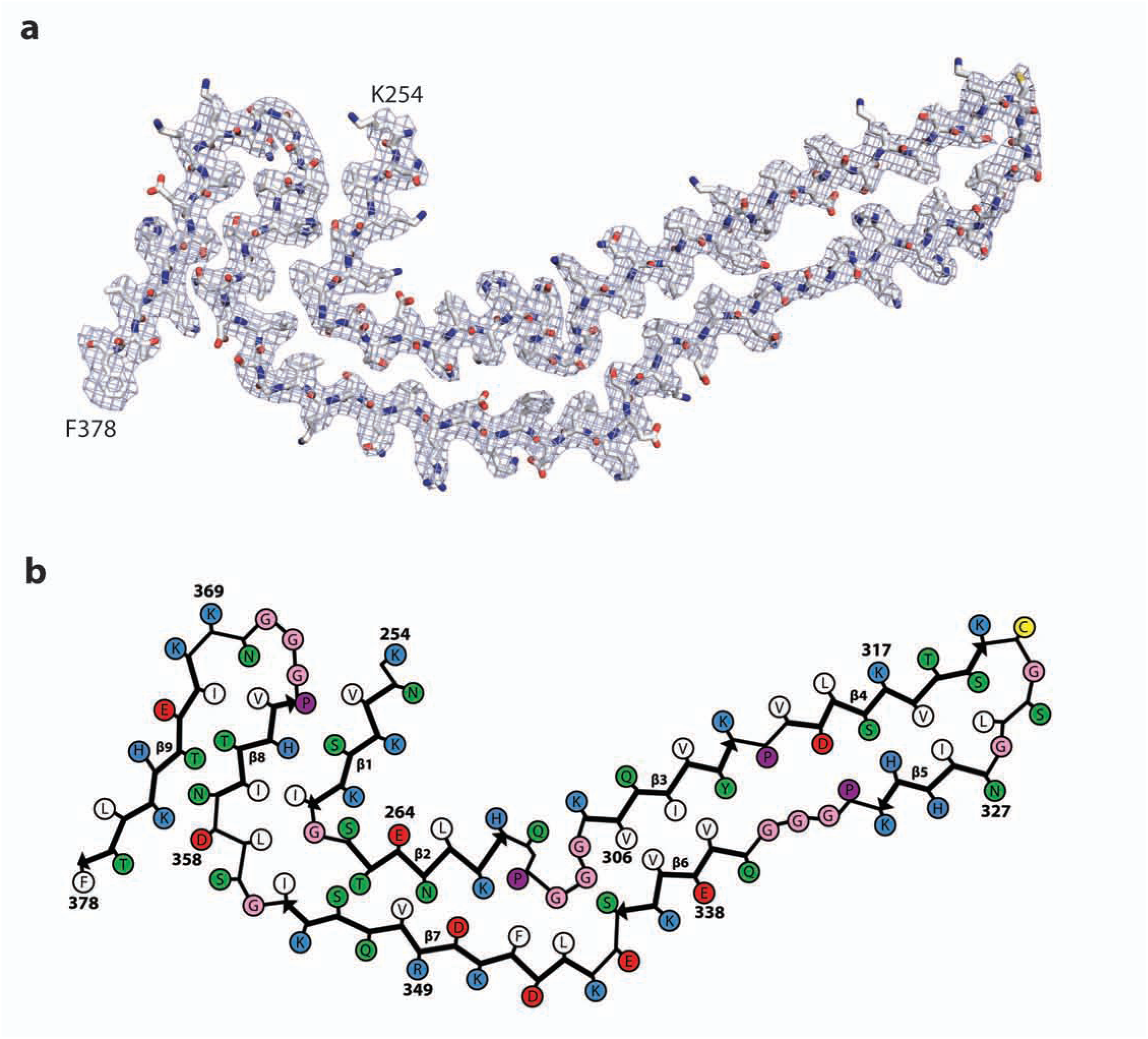
The Pick tau filament fold. **a.** Sharpened, high-resolution cryo-EM map of the narrow Pick filament (NPF) with the atomic model of the Pick fold overlayed. **b.** Schematic view of the Pick fold. Amino acid numbering corresponds to the 441 amino acid human tau isoform, so residues 275-305 of R2 are not present.

**Figure 3:**
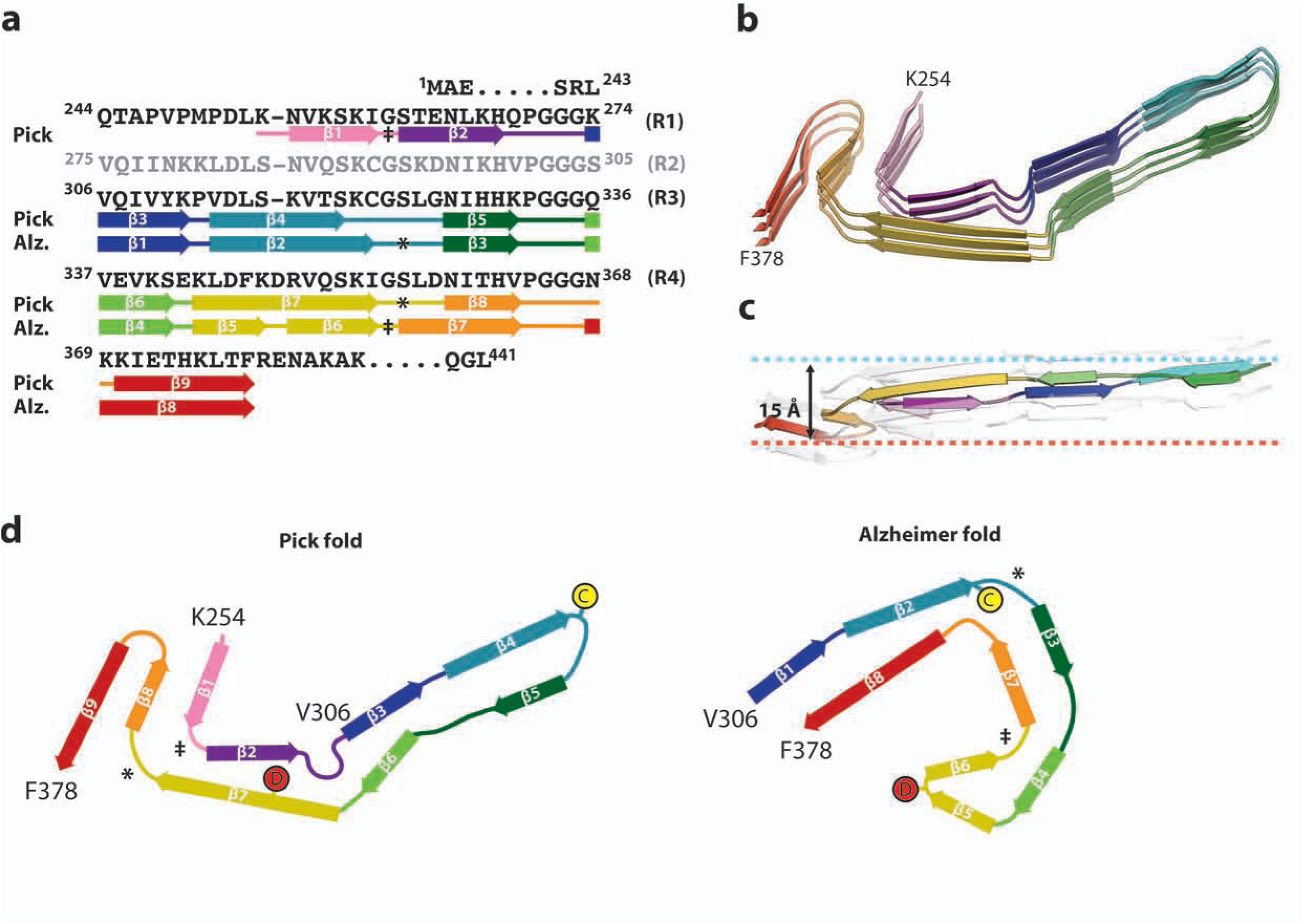
Comparison of the Pick and Alzheimer tau filament folds. a. Sequence alignment of the microtubule-binding repeats (R1–R4) with the observed nine (β-strand regions (arrows) in the Pick fold and eight (β-strand regions in the Alzheimer fold (arrows), coloured from violet to red. b. Rendered view of the secondary structure elements in the Pick fold, depicted as three successive rungs, c. As in b, but in a view perpendicular to the helical axis, revealing the changes in height within a single molecule, d. Schematic of the secondary structure elements in the Pick and Alzheimer folds, depicted as a single rung. The positions of C_322_ and D_348_ in the two folds are highlighted. The symbols ‡ and ∗ mark conserved turns of homologous regions in the Pick and Alzheimer folds.

The inter-strand connections and their interactions maintain the strand pairings and compensate for differences in strand lengths and orientations. A sharp right-angle turn at G_261_, between β1 and β2, faces a four-residue arc formed of _355_GSLD_358_, between *β*7 and *β*8, smoothly turning the chain direction at the same angle. The _270_PGGG_273_ motif between β2 and β3 forms an omega-shaped turn that compacts the protein chain locally, but maintains its direction at either end. On the opposite side, a β-arc formed of E_342_ and K_343_, between β6 and β7, creates space for this turn. In contrast, the homologous _332_PGGG_335_ motif connecting *β*5 and *β*6 forms an extended *β*-spiral conformation, compensating for the shorter lengths of these strands compared to the opposing β3 and β4, which are connected by P_312_. Solvent-mediated interactions may occur within the large cavity between this motif and the side-chains at the junction of β3 and β4. The third homologous _364_PGGG_367_ motif contributes to a 180° turn that allows *β*9 to pack against the other side of *β*8. Variations in the height of the chain along the helical axis also help to maintain an ordered hydrogen-bonding pattern of the β-stranded regions (Fig 3c).

The solvent-exposed side-chains of C_322_ and S_324_, together with the intervening G_323_, form a smooth flat surface at the hairpin-turn between β4 and β5. This provides the interface for the formation of WPFs by abutting of protofilaments (Extended Data Fig. 3). The distances between protofilaments at this interface would enable van der Waals interactions, but not disulfide bond formation. Stereochemically, domain-swapped tau dimers could also be accommodated within WPFs, whereby _322_CGSLG_326_ motifs would run antiparallel to each other, rather than forming hairpin turns, and the resulting interior C_322_ side-chains could form inter-chain disulfide bonds. However, the separation of the two protofilaments in the WPF reconstruction (Fig. 1g) and the observations that WPFs can lose segments of one protofilament and are stable under reducing conditions (Extended Data Fig. 3) lead us to conclude that WPFs are formed by two separate protofilaments making tight contacts at their distal tips through van der Waals interactions.

Three regions of less well-resolved density bordering the solvent-exposed faces of β4, β5 and β9 are apparent in the unsharpened maps of both NPFs and WPFs (Fig. 1f,g). Their low-resolution suggests that they represent less ordered, heterogeneous and/or transiently occupied structures. The density bordering β4 is similarly located, but more extended and less-well resolved, than that found to interact with the side-chains of K_317_, T_319_ and K_321_ in Alzheimer’s disease PHFs and SFs^11^, and hypothesized to be the N-terminal _7_EFE_9_, part of the discontinuous MC1 epitope^24^. NPFs and WPFs were labelled by MC1 (Extended Data Fig. 1).

It was not previously known why only 3R tau, which lacks the second microtubule-binding repeat, is present in Pick body filaments. Our results show that despite sequence homology, the structure formed by K_254_-K_274_ of the first tau repeat is inaccessible to the corresponding residues from the second repeat of 4R tau (S_285_-S_305_), because of the close packing between β2 and β7, which cannot accommodate the bulkier side-chain of K_294_ from 4R tau instead of T_263_ from 3R tau, and because the site preceding the omega-like structure formed by _270_PGGG_273_ cannot accommodate a Cβ branched residue, such as V_300_ from 4R tau instead of Q_269_ from 3R tau (Extended Data Fig. 4). In addition, the smaller C_291_ residue from 4R tau would form weaker interactions with L_357_ and I_360_ than those formed by I_260_ of 3R tau. In support, tau filaments extracted from the brain of the patient with Pick’s disease used for cryo-EM seeded the aggregation of recombinant full-length 3R, but not 4R, tau (Extended Data Fig. 5). Similar experiments have shown that Alzheimer’s disease PHFs and SFs, whose core sequences are shared by 3R and 4R tau, can seed both types of isoform^25^. Such templated misfolding explains the selective incorporation of 3R tau in Pick body filaments. Pick’s disease extracts have been reported to seed the aggregation of a 4R tau fragment comprising the repeats (residues 244-372) with mutations P301L and V337M^26^. However, this tau fragment cannot form the Pick fold, which is unable to accommodate R2 and requires residues 373-378. A small amount of aggregated four-repeat tau may have accounted for the seeding activity, as suggested in a separate study^8^. Loss of von Economo neurons in anterior cingulate and frontoinsular cortices has been reported to be an early event in Pick’s disease^27,28^. It remains to be established how 3R tau seeds can form in cells that also express 4R tau. Alternatively, nerve cell populations may be distinguished by the tau isoforms that they express^29^.

To test the generality of the Pick fold, we investigated the binding of repeat-specific antibodies to tau filaments extracted from the frontotemporal cortex of eight additional cases of sporadic Pick’s disease (Extended Data Table 1). By Western blotting, all samples ran as two tau bands of 60 and 64 kDa, which were detected by anti-R1, -R3 and -R4 antibodies, but not by an anti-R2 antibody, showing the presence of only 3R tau (Extended Data Fig. 6). Immunogold negative-stain electron microscopy showed that most filaments were NPFs, with a minority of WPFs, and were not decorated by the repeat-specific antibodies (Extended Data Fig. 7). This shows that the R1, R3 and R4 epitopes are inaccessible to the antibodies used, indicating that they form part of the ordered filament core. Alzheimer’s disease PHFs and SFs are decorated by anti-R1 and - R2, but not by anti-R3 and -R4 antibodies, because their core is made of R3, R4 and the 10 amino acids following R4^11,30^. These results are in good agreement with experiments using limited proteolysis and mass-spectrometry^7^. We conclude that the ordered core of tau filaments from Pick’s disease comprises the C-terminal part of R1, all of R3 and R4, as well as 10 amino acids after R4.

Unlike Alzheimer’s disease PHFs and SFs, Pick body filaments are not phosphorylated at S_262_ and/or S_356_ (Extended Data Fig. 1)^14,16^. The reasons for this differential phosphorylation are unknown. Our structure reveals that the tight turn at G_261_ prevents phosphorylation of S_262_ in the ordered core of Pick’s disease filaments, whereas the phosphorylated S_262_ is outside the ordered core of the Alzheimer tau filament fold^11^. This explains the differential phosphorylation and raises the question of whether phosphorylation at S_262_ may protect against Pick’s disease.

In the Pick and Alzheimer tau filament folds, most β-structure residues between V_306_ and I_354_ align locally, as do the connecting segments of P_312_, _332_PGGG_335_ and _342_EK_343_ (Figure 3a). Almost all amino acid side-chains from this region have the same interior or solvent-exposed orientations in both folds. Exceptions are C_322_ and D_348_, which cause reversed chain directions in one or other fold (Figure 3d). The side-chain of C_322_ is interior in the Alzheimer tau filament fold, whereas it is solvent-exposed in the Pick fold. This enables the hairpin-like turn and the cross-β packing of β4 against β5. The side-chain of D_348_ is interior in the Pick tau filament fold, thereby maintaining β-structure from K_343_ to I_354_ (β7), whereas it is solvent exposed in the Alzheimer fold, enabling the tight turn between β5 and β6, which, together with β4, gives rise to a triangular β-helix conformation^11^. Such β-helices, previously thought to be important for propagation^31^, are absent from the Pick tau filament fold. The β-strands in G_355_-F_378_ align well in both folds, but have different cross-β packing arrangements. The solvent-exposed side-chains of β7 and β8 in the Alzheimer fold are interior in the equivalent strands of the Pick fold (β8 and β9), because of different conformations of the two turn regions in R4, _355_GSLD_358_ and _364_PGGG_367_. The _355_GSLD_358_ motif makes a sharp right-angle turn at G_355_ in the Alzheimer tau filament fold, but a wide turn in the Pick fold. The same sharp turn is found at the homologous site in R1 in the Pick tau filament fold, whereas the same wide turn occurs at the homologous site in R3 in the Alzheimer fold (Fig. 3). This suggests that these semi-conserved turn structures may also be found in tau filament folds in other diseases. In contrast, the _364_PGGG_367_ motif adopts a new conformation in the Pick fold, which reverses the chain direction and is different from both the right-angle turn that this motif forms in the Alzheimer fold and the conformations of the homologous PGGG motifs from the other repeats in both tau filament folds. The Pick and Alzheimer folds share similar secondary structure patterns, but different turn conformations result in distinct cross-β packing.

These findings show that the ordered cores of tau filaments from Pick’s disease adopt a single, novel fold of 3R tau, which is distinct from the tau filament fold of Alzheimer’s disease. This suggests that different folds may account for tauopathies with 4R tau filaments, such as progressive supranuclear palsy. Our results also suggest that single, disease-specific folds may exist in tauopathies with the same tau filament isoform composition, such as progressive supranuclear palsy and corticobasal degeneration, since identical tau sequences can adopt more than one fold. Conserved secondary structure motifs and markedly different conformations at turn residues in the Alzheimer and Pick tau filament folds may form the basis for structural diversity in tau protein folds from other neurodegenerative diseases.

The identification of disease-specific folds in the ordered cores of tau filaments establishes the existence of molecular conformers. This is central to the hypothesis that conformers of filamentous tau give rise to the clinical phenotypes that define distinct tauopathies, akin to prion strains. By revealing the structural basis for molecular conformers in specific diseases, our results pave the way to a better understanding of a wide range of diseases related to abnormal protein aggregation.

## Acknowledgements

We thank the patients’ families for donating brain tissue; M. R. Farlow for clinical evaluation; F. Epperson, R. M. Richardson and U. Kuederli for human brain collection and analysis; P. Davies, P. Seubert and M. Hasegawa for antibodies MC-1, 12E8 and TauC4, respectively; S. Chen, C. Savva and G. Cannone for support with electron microscopy; T. Darling and J. Grimmett for help with computing; W.W. Seeley and M.G. Spillantini for helpful discussions. M.G. is an Honorary Professor in the Department of Clinical Neurosciences of the University of Cambridge. This work was supported by the UK Medical Research Council (MC_UP_A025_1012 to G.M., MC_UP_A025_1013 to S.H.W.S. and MC_U105184291 to M.G.), the European Union (Joint Programme-Neurodegeneration Research REfrAME to M.G. and B.F. and the Innovative Medicines Initiative 2 IMPRiND, project number 115881, to M.G.), the US National Institutes of Health (grant P30-AG010133 to B.G.), the Department of Pathology and Laboratory Medicine, Indiana University School of Medicine (to B.G.) and the Alzheimer’s Association Zenith Award (to R.V.).

## Contributions

B.G. performed neuropathology; H.J.G. and R.V. carried out genetic analysis; B.F. extracted tau filaments; B.F. and W.Z. conducted immunolabelling; B.F. and W.Z. purified recombinant tau proteins; B.F. carried out seeded aggregation; B.F. and W.Z. performed cryo-EM; B.F., W.Z. and S.H.W.S. analysed the cryo-EM data; B.F., W.Z., G.M. and A.M. built the atomic model; R.A.C. contributed to the inception of the study; M.G. and S.H.W.S. supervised the project; all authors contributed to writing the manuscript.

## Methods

### Extraction of tau filaments

Sarkosyl-insoluble material was extracted from grey matter of frontal and temporal cortex of the patients’ brains, as described^1^. The pelleted sarkosyl-insoluble material was resuspended in 50 mM Tris-HCl pH 7.4 containing 150 mM NaCl and 0.02% amphipol A8-35 at 250 μl per g tissue, followed by centrifugation at 3,000 x*g* for 30 min at 4 °C. The pellets, containing large contaminants, were discarded. The supernatants were centrifuged at 100,000 x*g* for 30 min at 4 °C. The resulting pellets were resuspended in buffer at 15 μl per g tissue. Pronase treatment was carried out as described for negative-stain EM^1^ and cryo-EM^2^.

### Cloning and purification of epitope-deletion recombinant tau

Tau constructs lacking the BR136, Anti-4R, BR135 and TauC4 peptide sequences were cloned from pRK172 encoding wild-type 0N4R or 2N4R tau using the QuikChange Lightning site-directed mutagenesis kit (Agilent), according to the manufacturer’s instructions. Recombinant proteins were purified as described^3^.

### Immunolabelling and histology

Western blotting and immunogold negative-stain EM were carried out as described^1^. For Western blotting, samples were resolved on 4–20% or 10% Tris-glycine gels (Novex), and the primary antibodies were diluted in PBS plus 0.1% Tween 20 and 1% BSA. BR136 is a polyclonal antibody that was raised against a synthetic peptide corresponding to residues 244-257 of tau. The peptide (200 μg), coupled to keyhole limpet hemocyanin using glutaraldehyde, was mixed 1:1 with Freund’s complete adjuvant and used to immunise white Dutch rabbits. Booster injections using 200 μg of conjugated peptide mixed 1:1 with Freund’s incomplete adjuvant were given every 2 weeks for 10 weeks following the primary immunisation. Antibodies were harvested 7 days after the final booster injection and affinity purified. Extended Data Figure 6 shows that BR136 is specific for the C-terminal region of residues 244-257. Neurohistology and immunohistochemistry were carried out as described^4^. Brain sections were 8 μm thick and were counterstained with haematoxylin. Detailed antibody information is provided in Extended Data Table 2.

### Whole exome sequencing

Whole exome sequencing was carried out at the Center for Medical Genomics of Indiana University School of Medicine using genomic DNA from the nine individuals with neuropathologically confirmed diagnoses of Pick’s disease, the tau filaments of which were used in Extended Data Figures 6 and 7. Target enrichment was performed using the SureSelectXT human all exon library (V6, 58Mb, Agilent) and high-throughput sequencing using a HiSeq4000 (2×75bp paired-end configuration, Illumina). Bioinformatics analyses were performed as described^5^. Findings on *MAPT*, *PSEN1* and *APOE* are presented in Extended Data Table 1.

### Seeded aggregation

Seeded aggregation was carried out as described^6^, but with full-length wild-type tau protein and without the aggregation inducer heparin. Recombinant 0N3R and 0N4R tau were purified as described^3^. Extracted tau filaments (15 μl per g tissue) were diluted 1:10 in 10 mM HEPES pH7.4, 200 mM NaCl; 2 μl was added to 98 μl of 20 μM 0N3R or 0N4R recombinant tau in the same buffer with 10 μM thioflavin T in a black, clear bottom 96-well plate (Perkin Elmer). The plate was sealed and incubated at 37 °C in a plate reader (BMG Labtech FLUOstar Omega), with cycles of shaking for 60s (500 rpm, orbital) followed by no shaking for 60 s. Filament formation was monitored by measuring Thioflavin T fluorescence every 45 min using 450 −10 nm excitation and 480 −10 nm emission wavelengths, with an instrument gain of 1100. Three independent experiments were performed with separate recombinant protein preparations.

### Electron cryo-microscopy

Extracted, pronase-treated tau filaments (3 μl at a concentration of ~0.5 mg/ml) were applied to glow-discharged holey carbon grids (Quantifoil Au R1.2/1.3, 300 mesh) and plunge-frozen in liquid ethane using an FEI Vitrobot Mark IV. Images were acquired on a Gatan K2-Summit detector in counting mode using an FEI Titan Krios at 300 kV. A GIF-quantum energy filter (Gatan) was used with a slit width of 20 eV to remove inelastically scattered electrons. Fifty-two movie frames were recorded, each with an exposure time of 250 ms using a dose rate of 1.06 electrons per Å^2^ per frame for a total accumulated dose of 55 electrons per Å^2^ at a pixel size of 1.15 Å on the specimen. Defocus values ranged from –1.7 to –2.8 μm. Further details are presented in Exended Data Table 3.

### Helical reconstruction

Movie frames were corrected for gain reference, motion-corrected and dose-weighted using MOTIONCOR2^7^. Aligned, non-dose-weighted micrographs were used to estimate the contrast transfer function (CTF) in Gctf^8^. All subsequent image-processing steps were performed using helical reconstruction methods in RELION 2.1^9,10^. NPFs and WPFs were picked manually and processed as separate datasets.

#### NPF dataset

NPF segments were extracted using a box size of 270 pixels and an inter-box distance of ~10% of the box size. Reference-free 2D classification was performed using a regularization value of T = 2, and segments contributing to suboptimal 2D class averages were discarded. An initial helical twist of –0.73° was estimated from the crossover distances of NPFs in micrographs, and the helical rise was estimated to be 4.7 Å. Using these values, an initial 3D reference was reconstructed *de novo* from 2D class averages of segments comprising an entire helical cross-over. A first round of 3D classification, starting from the *de novo* initial model low-pass filtered to 40 Å, with local optimization of the helical twist and rise, and a regularization value of T = 4 yielded a reconstruction in which individual β-sheets perpendicular to the helical axis were clearly separated, but no structure was discernable along the helical axis. Subsequently, 3D auto-refinement with optimization of helical twist and rise and a regularization value of T = 10 was performed using the segments that contributed to the 3D class displaying β-sheets. The resulting reconstruction showed clearly discernable β-strand separation.

An additional round of 3D classification with a regularization value of T = 10 starting from the 5 Å low-pass filtered map from the previous auto-refinement was used to further select segments for a final high-resolution refinement. In total, 16,097 segments contributed to the final map. The reconstruction obtained with this relatively small subset of the initial dataset matched lower-resolution reconstructions obtained with larger subsets of the data, indicating that image classification did not select for a specific structure from a conformationally heterogeneous dataset, but instead was successful in distinguishing the segments with high-resolution information from images of varying quality. This is in line with observations in single-particle analysis^11^. Superimposing the selected segments onto the original micrographs further confirmed this. Image classification also did not separate filaments with variable twists; instead, RELION combines segments from filaments with variable twists into a single 3D reconstruction and reduces the corresponding blurring effects by only using the central part of an intermediate asymmetrical reconstruction for real-space helical symmetrisation^10^. We used a 10% value for the corresponding helical z_percentage parameter.

Optimization of the helical twist and rise converged onto –0.75° and 4.78 Å, respectively. Refinements with helical rises of multiples of 4.78 Å all led to β-strand separation, but in agreement with the observed absence of layer lines between 50 and 4.7 Å we were unable to detect any repeating patterns along the helical axis other than the successive rungs of β-strands.

The final NPF reconstruction was sharpened using standard post-processing procedures in RELION, resulting in a B-factor of –57 Å^2^ (Extended Data Table 2). Helical symmetry was imposed on the post-processed map using RELION helix toolbox^10^. Final, overall resolution estimates were calculated from Fourier shell correlations at 0.143 between the two independently refined half-maps, using phase-randomization to correct for convolution effects of a generous, soft-edged solvent mask^12^. The overall resolution estimate of the final map was 3.2 Å. Local resolution estimates were obtained using the same phase-randomization procedure, but with a soft spherical mask that was moved over the entire map.

#### WPF dataset

The WPF dataset was down-scaled to a pixel size of 3.45 Å and segments were extracted using a box size of 180 pixels and an inter-box distance of ~10% of the box size. As with the NPF dataset, an initial 3D reference was reconstructed *de novo* from 2D class averages of segments comprising an entire helical cross-over. 3D classification was then performed to discard suboptimal segments. 3D auto-refinement of the best class with a regularization value of T = 4 and a fixed helical rise and twist of 4.7 Å and –0.6°, respectively, led to a 3D structure with good separation of β-sheets perpendicular to the helical axis, but no structure was discernable along the helical axis. The cross-section of this map clearly revealed the presence of two NPF protofilaments. To further improve the reconstruction, we also made an initial model by placing two NPF maps, rotated 180° relative to each other in the WPF reconstruction, and low-pass filtering the resulting map to 60 Å. After a second 3D auto-refinement starting from this model, the final WPF reconstruction had an estimated overall resolution of 8 Å and was sharpened by specifying a b-factor of −200 Å^2^ (Extended Data Table 2).

In total, 3,003 segments contributed to the final map.

### Model building and refinement

A single monomer of the NPF core was built *de novo* in the 3.2 Å resolution reconstruction using COOT^13^. Model building was started from the distinctive extended *β*-spiral conformation of the _332_PGGG_335_ motif, neighbouring the large histidine side chains of residues 329 and 330, and working towards the N- and C-terminal regions by manually adding amino acids, followed by targeted real-space refinement. This model was then translated to give a stack of three consecutive monomers to preserve nearest-neighbour interactions for the middle chain in subsequent refinements using a combination of real-space refinement in PHENIX^14^ and Fourier-space refinement in REFMAC^15^. In the latter, local symmetry restraints were imposed to keep all *β*-strand rungs identical. Since most of the structure adopts a *β*-strand conformation, hydrogen-bond restraints were imposed to preserve a parallel, in-register hydrogen-bonding pattern in earlier stages of the model building process. Side-chain clashes were detected using MOLPROBITY^16^ and corrected by iterative cycles of real-space refinement in COOT and Fourier-space refinement in REFMAC. The refined model of the NPF was rigid-body fitted into the WPF map. Separate NPF model refinements were performed against a single half-map, and the resulting model was compared to the other half-map to confirm the absence of overfitting. The final model was stable in refinements without additional restraints.

### Ethical review board and informed consent

The Indiana Alzheimer Disease Center studies were reviewed and approved by the Indiana University Institutional Review Board. Informed consent was obtained from the patients’ next of kin.

### Data availability

Cryo-EM maps and the refined atomic model will be deposited in the Electron Microscopy Data Bank and the Protein Data Bank, respectively, following acceptance of the manuscript.

---

**Extended Data Figure 1:**
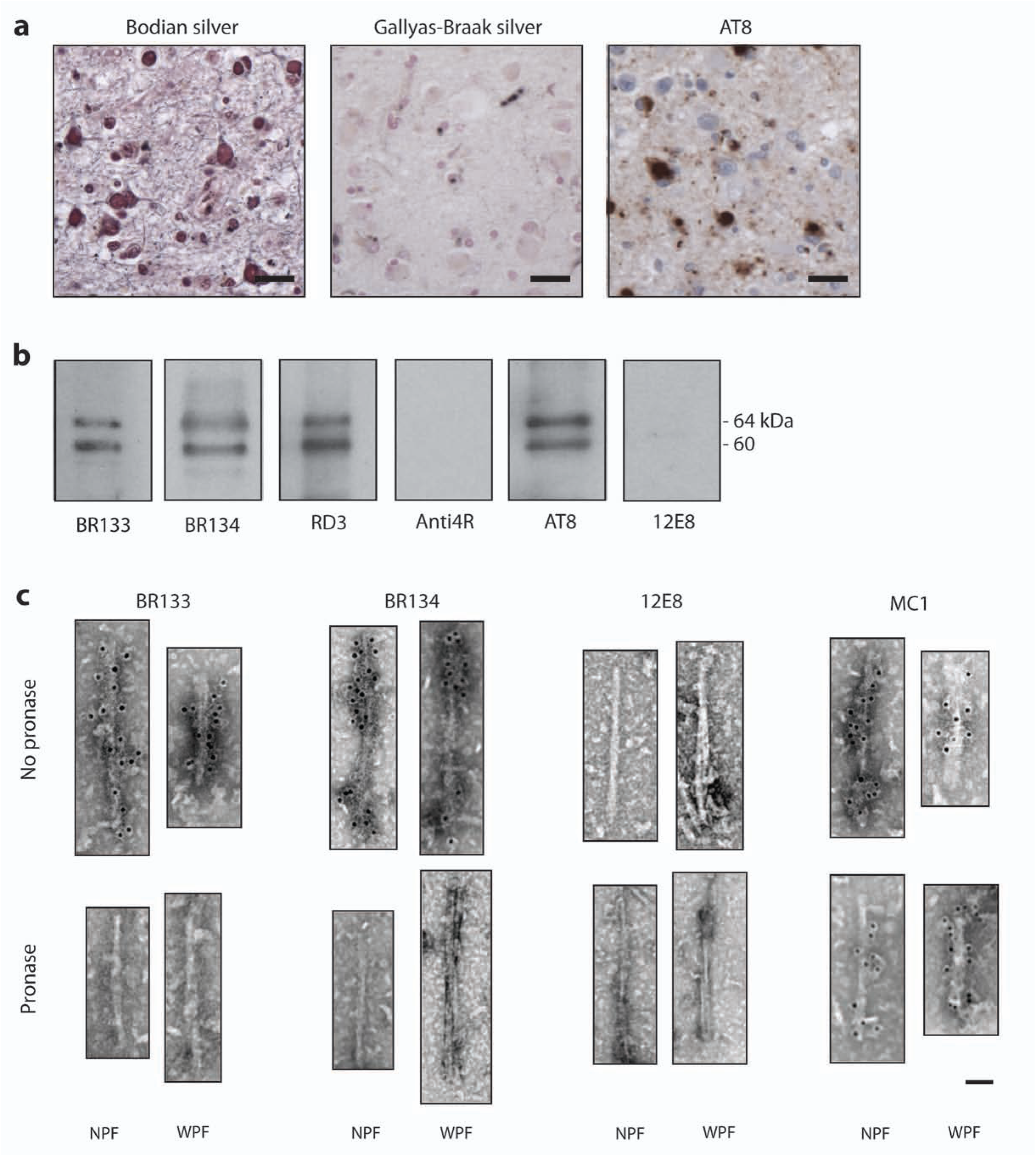
Further characterisation of the filamentous tau pathology of Pick’s disease. **a.** Light microscopy of sections from the frontotemporal cortex showing staining of Pick bodies using Bodian silver and antibody AT8, but not Gallyas-Braak silver. Nuclei are counterstained blue. Scale bars, 50 μm **b,c.** Immunolabeling of the sarkosyl-insoluble fraction from the patient’s frontotemporal cortex. Immunoblots (**b**) using anti-Tau antibodies BRI 33 (amino-terminus), BRI 34 (carboxy-terminus), RD3 (3R tau), Anti-4R (4R tau), AT8 (pS202 and pT205) and 12E8 (pS262 and/or pS356). Immunogold negative-stain electron microscopy (**c**) of NPFs and WPFs with BR133, BR134,12E8 and MC1 with and without mild pronase treatment. Scale bar, 500 Å.

**Extended Data Figure 2:**
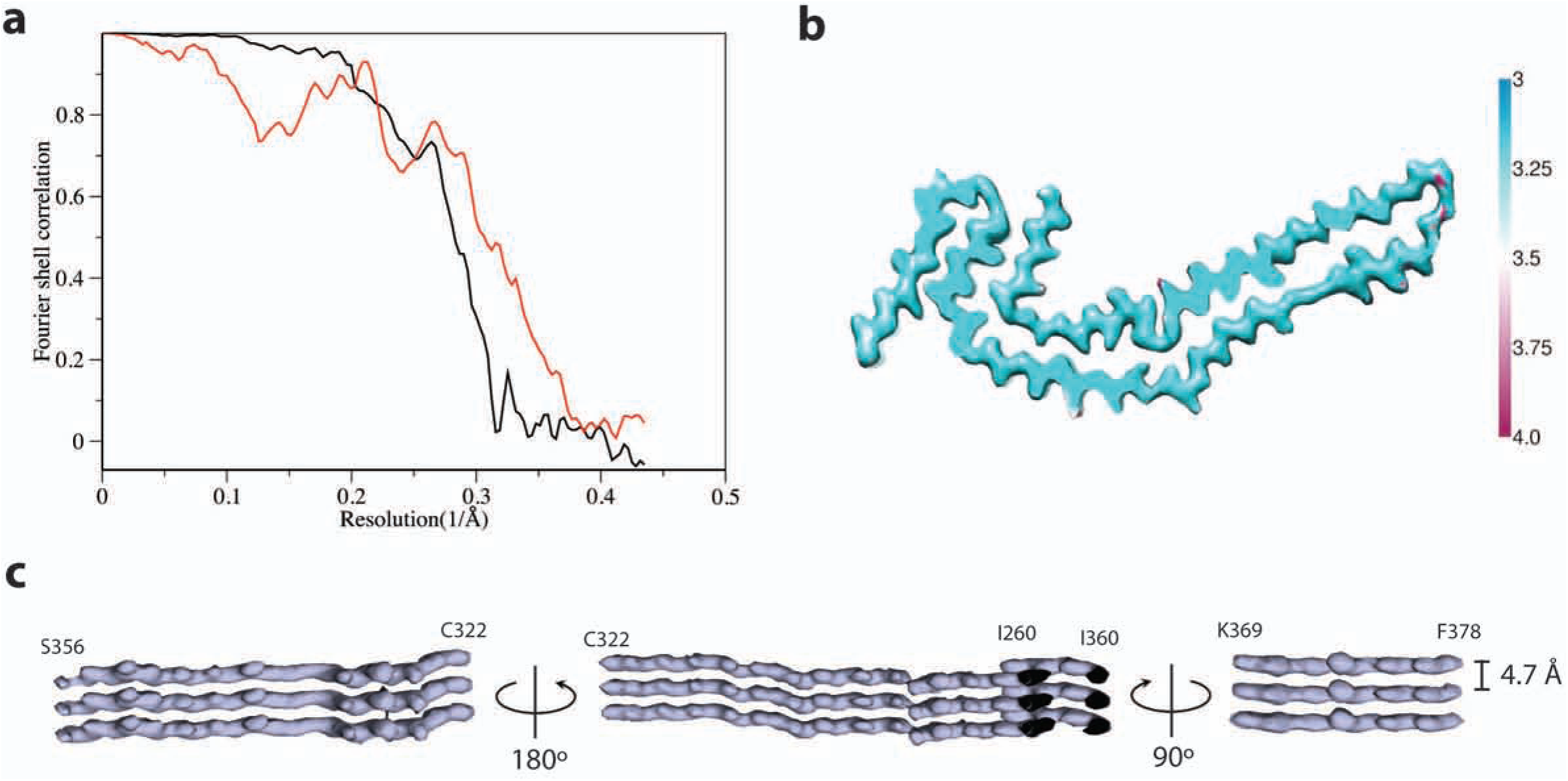
Narrow Pick filament (NPF) structure. **a.** Fourier shell correlation curves between two independently refined half-maps (black line) and between the cryo-EM reconstruction and refined atomic model (red line), **b.** Local resolution estimates for the NPF reconstruction, **c.** Helical axis views of the NPF reconstruction.

**Extended Data Figure 3:**
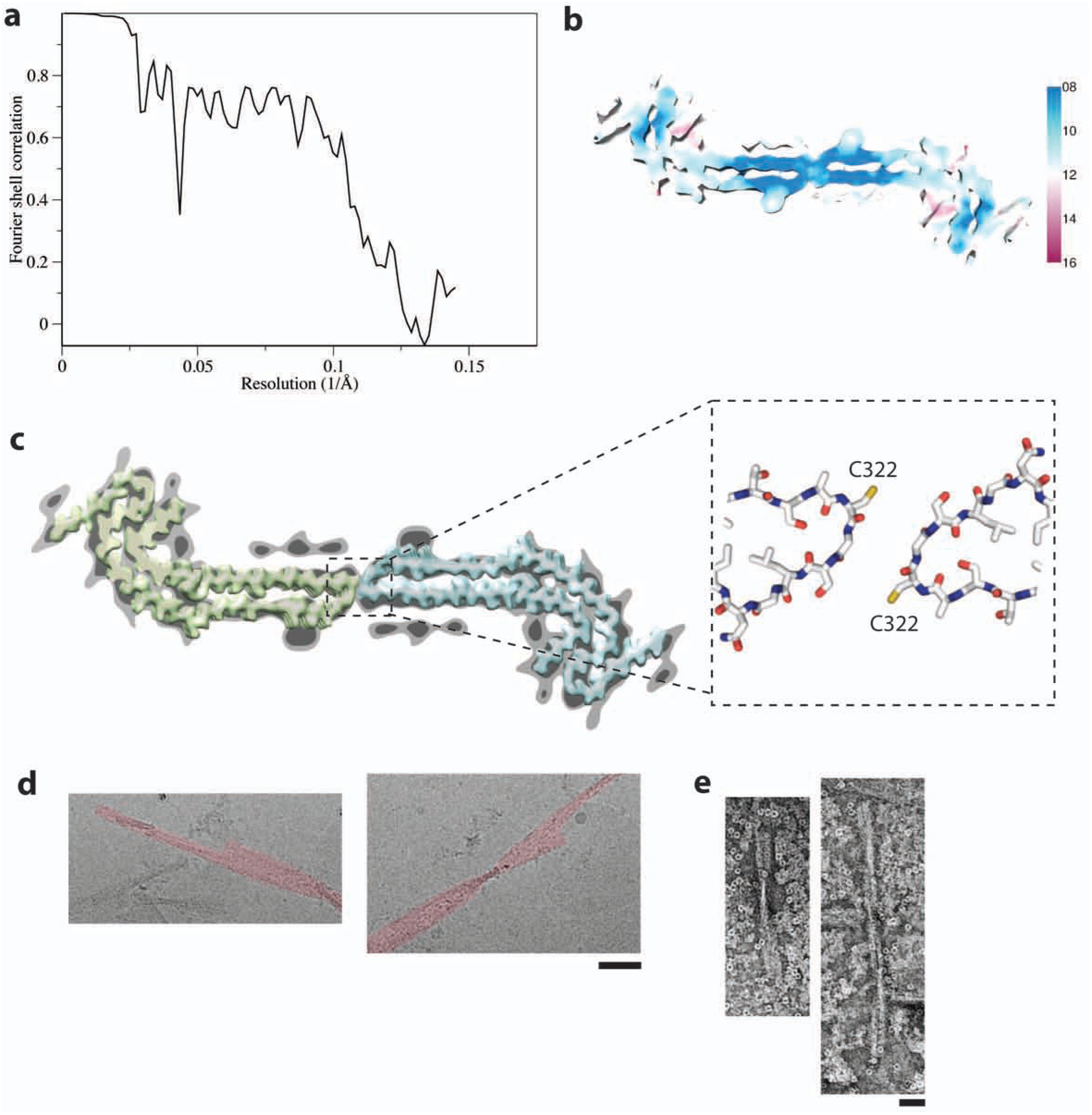
Wide Pick filament (WPF) structure. **a.** Fourier shell correlation curves between two independently refined half-maps. **b.** Local resolution estimates for the WPF reconstruction, **c.** WPF density at high (light grey) and low (dark grey) threshold with densities for two NPFs overlaid (yellow and blue). The atomic models fitted to the NPF densities in the region of the protofilament interface are shown in the boxed out area. **d.** Cryo-EM images showing WPFs (false coloured red) where segments from one of the protofilaments have been lost. Scale bar, 500 Å. **e.** Negative-stain EM images of WPFs following incubation in 100 mM dithiothreitol for 20 h. Scale bar, 500 Å.

**Extended Data Figure 4:**
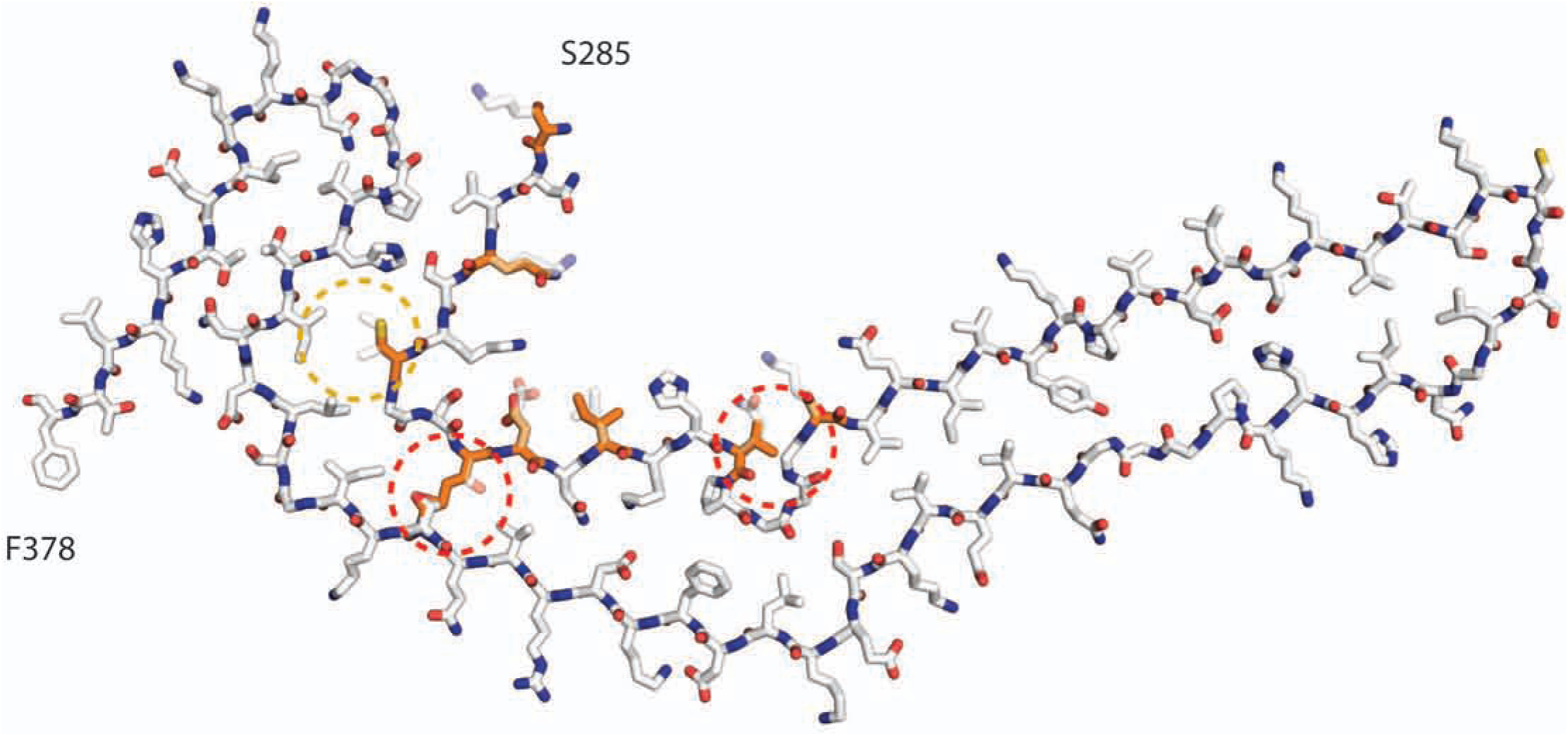
Incompatibility of Pick tau filament fold with 4R tau. Atomic model of Pick fold with 4R tau sequence overlaid. The region formed by K254-K274 from R1 is replaced by the S285-V300 region from R2 in 4R tau. Residues that differ between these regions of R1 and R2 are coloured orange. The major discrepancies of lysine at position 294 in R2, instead of threonine at position 263 in R1, and valine at position 300 in R2, instead of glutamine at position 269 in R1, are highlighted with dashed red outlines. The minor discrepancy of weaker interactions of C291 of R2 with L357 and I360 than those formed by I260 of R1 is highlighted with a dashed yellow outline.

**Extended Data Figure 5:**
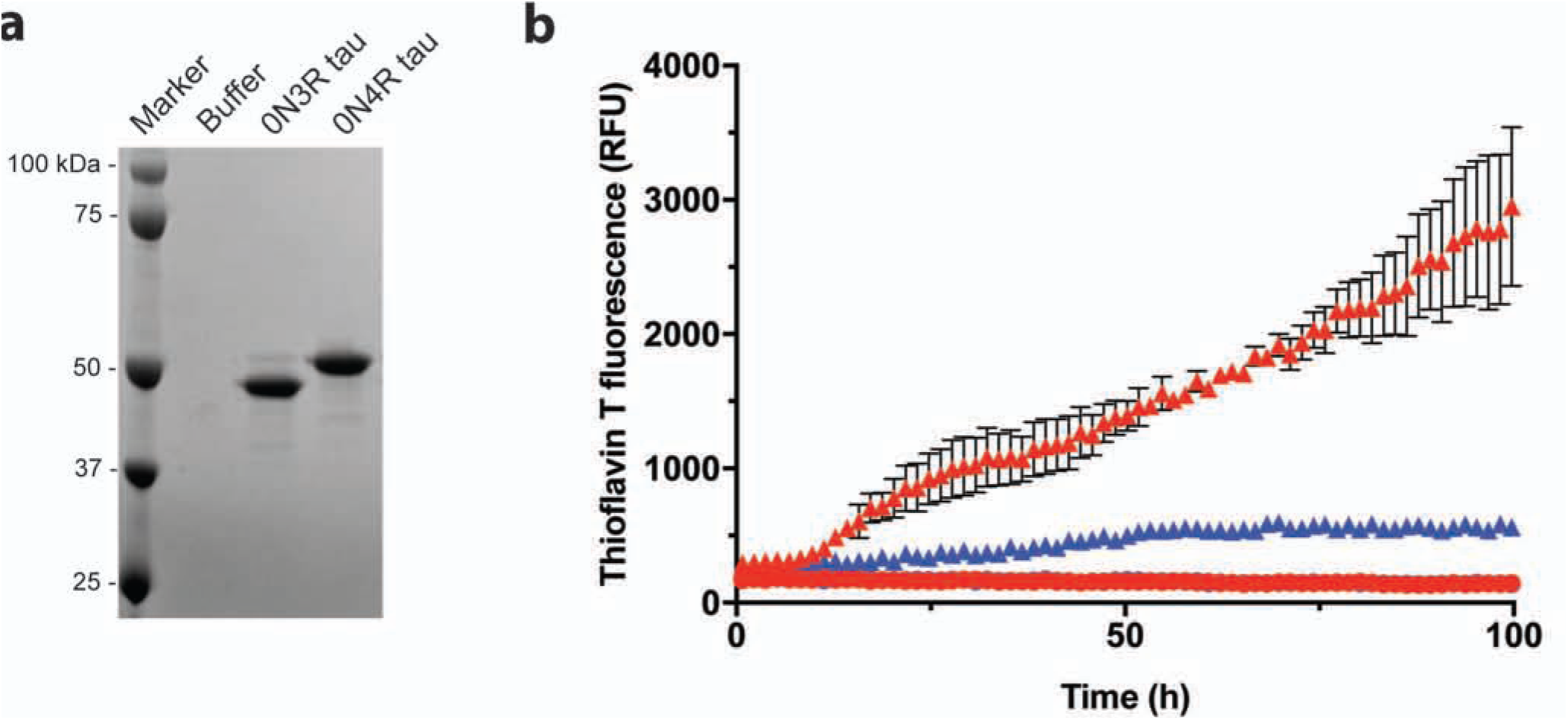
Seeded aggregation of full-length 3R, but not 4R, tau by the sarkosyl-insoluble fraction from Pick’s disease brain. **a.** Coomassie-stained SDS-PAGE of the substrates used for seeded aggregation, **b.** Thioflavin T fluorescence measurements of 0N3R (red) and 0N4R (blue) recombinant tau following incubation with (triangles) or without (circles) the sarkosyl-insoluble fraction from Pick’s disease brain used for cryo-EM. The results are expressed as the means ± SEM of three independent experiments using separate recombinant protein preparations. Error bars shorter than data point symbols are not shown. The sarkosyl-insoluble fraction from Pick’s disease brain efficiently seeded the aggregation of 3R, but not 4R, tau.

**Extended Data Figure 6:**
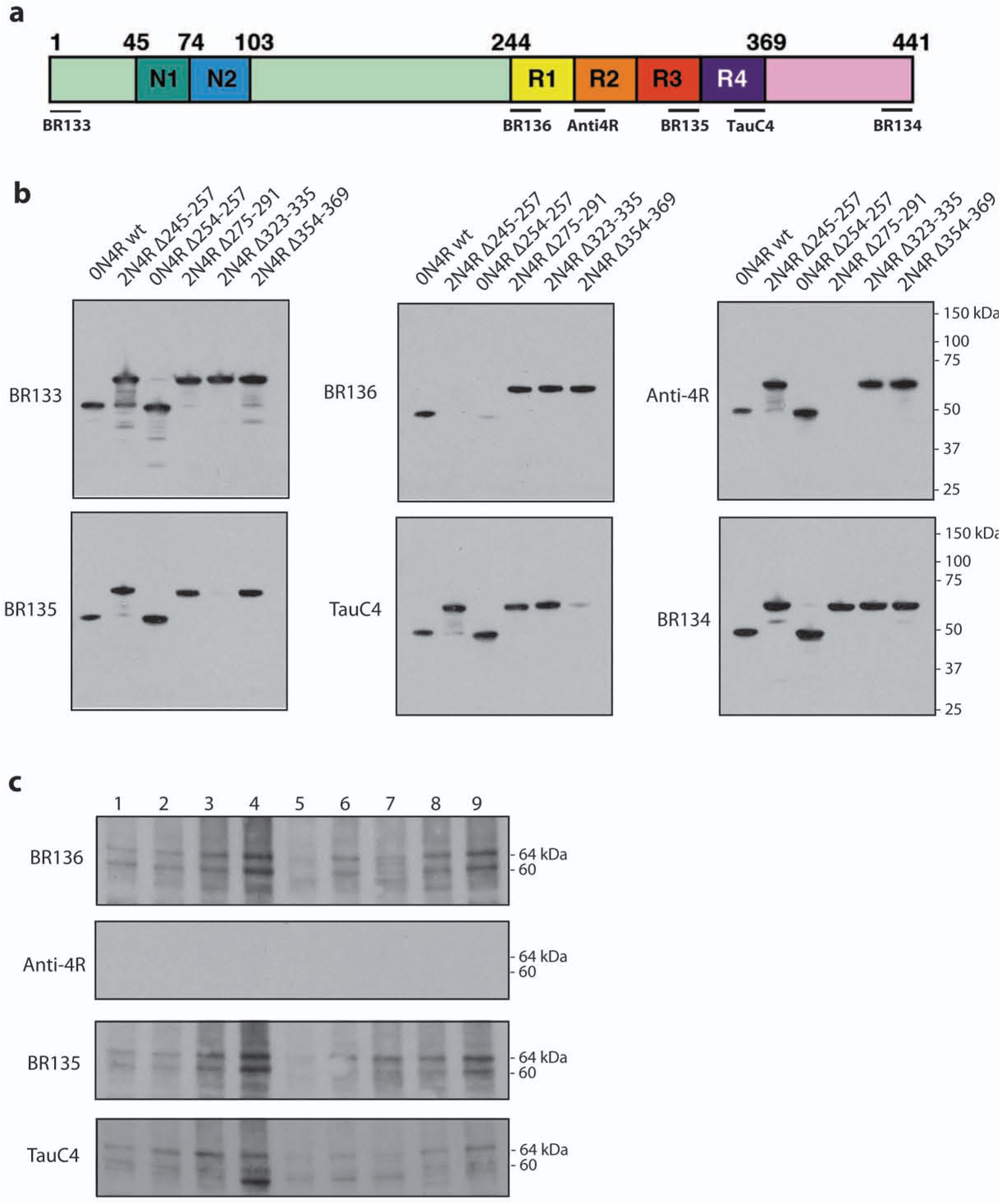
Immunoblot analysis of additional Pick’s disease cases. **a.** Diagram of 2N4R tau showing the N-terminal inserts (N1, N2), the repeats (R1-R4) and the epitopes of antibodies BRI 33 (N-terminus), BRI 36 (R1), Anti-4R (R2), BRI 35 (R3), TauC4 (R4) and BR134 (C-terminus). **b.** Immunoblots of epitope-deletion recombinant tau constructs with the antibodies shown in a. **c.** Immunoblots using the antibodies BR136, Anti-4R, BR135 andTauC4 of tau filaments extracted from frontotemporal cortex of 9 cases of Pick’s disease.

**Extended Data Figure 7:**
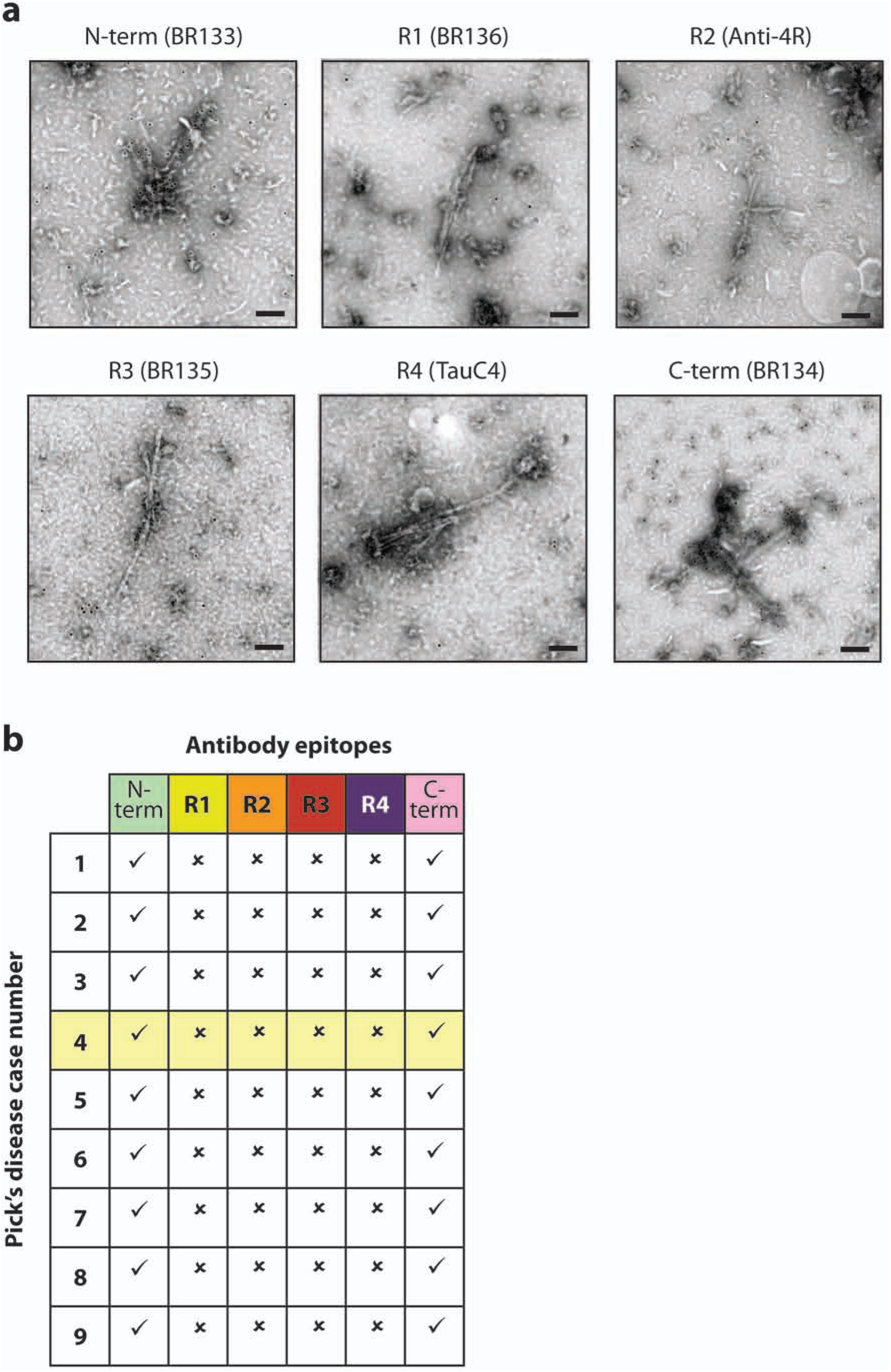
Immunogold negative-stain EM analysis of additional Pick’s disease cases. **a.** Representative immunogold negative-stain electron microscopy of NPFs and WPFs extracted from frontotemporal cortex of Pick’s disease brain (case number 4, which was also used for cryo-EM, highlighted in yellow) with antibodies against tau N-terminus (BR133), R1 (BR136), R2 (Anti4R), R3 (BR135), R4 (TauC4) and C-terminus (BR134). Scale bars, 100 nm. **b.** Table summarizing results from immunogold negative-stain electron microscopy of NPFs and WPFs extracted from frontotemporal cortex of 9 cases of Pick’s disease, as in a. See Extended Data Table 1 for details of Pick’s disease cases. Tick marks indicate antibody decoration of filaments, while crosses indicate that the antibodies did not decorate filaments. NPFs and WPFs were decorated by the antibodies against the N- and C-termini, but not by the repeat-specific antibodies.

**Extended Data Table 1:** Summary of Pick’s disease patients. Wild-type (wt) means that no known disease-causing mutations in the tau gene (*MAPT*) or the presenilin-1 gene (*PSEN1*) were detected. The patient used for cryo-EM is highlighted in yellow.

| Patient | Gender | Age at death (years) | *MAPT* | *PSEN1* | *APOE* haplotypes | Post-mortem interval (h) |
| --- | --- | --- | --- | --- | --- | --- |
| 1 | F | 73 | wt | wt | ε3/ε3 | 23.8 |
| 2 | F | 70 | wt | wt | ε3/ε3 | 20.5 |
| 3 | M | 61 | wt | wt | ε3/ε4 | 24.5 |
| 4 | F | 63 | wt | wt | ε2/ε3 | 3.0 |
| 5 | M | 70 | wt | wt | ε3/ε3 | 14.0 |
| 6 | M | 65 | wt | wt | ε3/ε3 | 12.5 |
| 7 | M | 64 | wt | wt | ε3/ε4 | 3.0 |
| 8 | M | 56 | wt | wt | ε3/ε4 | 9.0 |
| 9 | M | 69 | wt | wt | ε3/ε4 | 4.5 |

**Extended Data Table 2:**
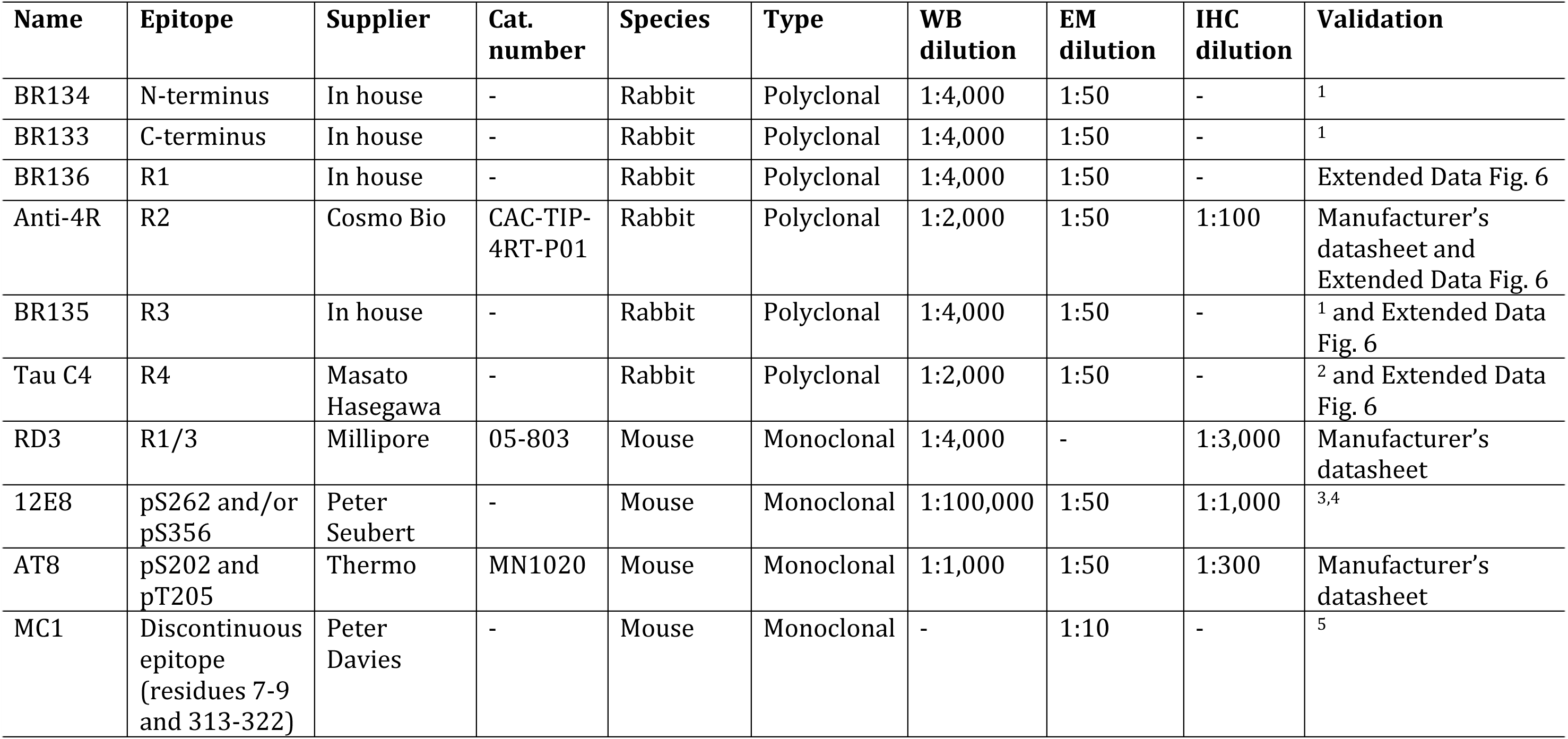
Primary anti-tau antibodies used in this study. WB, Western blot; EM, Immunogold negative stain electron microscopy; IHC, Immunohistochemistry 1. Goedert, M., Spillantini, M. G., Jakes, R., Rutherford, D. & Crowther, R. A. Multiple isoforms of human microtubule-associated protein tau: sequences and localization in neurofibrillary tangles of Alzheimer’s disease. *Neuron* **3**, 519-526 (1989).
2. Taniguchi-Watanabe, S. *et al.* Biochemical classification of tauopathies by immunoblot, protein sequence and mass spectrometric analyses of sarkosyl-insoluble and trypsin-resistant tau. *Acta Neuropathol.* **131**, 267-280 (2016).
3. Seubert, P. *et al.* Detection of phosphorylated Ser262 in fetal tau, adult tau, and paired helical filament tau. *J. Biol. Chem.* **270**, 18917-18922 (1995).
4. Litersky, J. M. *et al.* Tau protein is phosphorylated by cyclic AMP-dependent protein kinase and calcium/calmodulin-dependent protein kinase II within its microtubule-binding domains at Ser-262 and Ser-356. *Biochem. J.* **316 (Pt 2)**, 655-660 (1996).
5. Jicha, G. A., Bowser, R., Kazam, I. G. & Davies, P. Alz-50 and MC-1, a new monoclonal antibody raised to paired helical filaments, recognize conformational epitopes on recombinant tau. *J. Neurosci. Res.* **48**, 128-132 (1997).

| Name | Epitope | Supplier | Cat. number | Species | Type | WB dilution | EM dilution | IHC dilution | Validation |
| --- | --- | --- | --- | --- | --- | --- | --- | --- | --- |
| BR134 | N-terminus | In house | - | Rabbit | Polyclonal | 1:4,000 | 1:50 | - | <sup>1</sup> |
| BR133 | C-terminus | In house | - | Rabbit | Polyclonal | 1:4,000 | 1:50 | - | <sup>1</sup> |
| BR136 | R1 | In house | - | Rabbit | Polyclonal | 1:4,000 | 1:50 | - | Extended Data Fig. 6 |
| Anti-4R | R2 | Cosmo Bio | CAC-TIP-4RT-P01 | Rabbit | Polyclonal | 1:2,000 | 1:50 | 1:100 | Manufacturer's datasheet and Extended Data Fig. 6 |
| BR135 | R3 | In house | - | Rabbit | Polyclonal | 1:4,000 | 1:50 | - | <sup>1</sup> and Extended Data Fig. 6 |
| Tau C4 | R4 | Masato Hasegawa | - | Rabbit | Polyclonal | 1:2,000 | 1:50 | - | <sup>2</sup> and Extended Data Fig. 6 |
| RD3 | R1/3 | Millipore | 05-803 | Mouse | Monoclonal | 1:4,000 | - | 1:3,000 | Manufacturer's datasheet |
| 12E8 | pS262 and/or pS356 | Peter Seubert | - | Mouse | Monoclonal | 1:100,000 | 1:50 | 1:1,000 | <sup>3,4</sup> |
| AT8 | pS202 and pT205 | Thermo | MN1020 | Mouse | Monoclonal | 1:1,000 | 1:50 | 1:300 | Manufacturer's datasheet |
| MC1 | Discontinuous epitope (residues 7-9 and 313-322) | Peter Davies | - | Mouse | Monoclonal | - | 1:10 | - | <sup>5</sup> |

**Extended Data Table 3:** Cryo-electron microscopy structure determination.

| **Data Collection** | **NPF** | **PHF** |
| --- | --- | --- |
| Magnification | x105,000 | x105,000 |
| Defocus range (μm) | -1.7 to -2.8 | -1.7 to -2.8 |
| Voltage (kV) | 300 | 300 |
| Microscope | Titan Krios | Titan Krios |
| Detector | K2 Summit | K2 Summit |
| Frame exposure time (ms) | 250 | 250 |
| Number of frames | 52 | 52 |
| Total dose (e<sup>-</sup>/Å<sup>-2</sup>) | 55 | 55 |
| Pixel size (Å) | 1.15 | 1.15 (downscaled 3.45) |
| **Reconstruction** |  |  |
| Box size (pixel) | 270 | 180 |
| Inter-box distance (Å) | 28 | 18 |
| Segments extracted | 83,475 | 8,024 |
| Segments after Class2D | 46,305 | - |
| Segments after Class3D | 16,097 | 3,003 |
| Resolution (Å) | 3.2 | 8 |
| B-factor (Å<sup>2</sup>) | -57 | -200 |
| Helical Rise (Å) | 4.78 | 4.7 |
| Helical twist (°) | -0.75 | -0.6 |

